# Composite modeling of leaf shape across shoots discriminates *Vitis* species better than individual leaves

**DOI:** 10.1101/2020.06.22.163899

**Authors:** Abigail E. Bryson, Maya Wilson Brown, Joey Mullins, Wei Dong, Keivan Bahmani, Nolan Bornowski, Christina Chiu, Philip Engelgau, Bethany Gettings, Fabio Gomezcano, Luke M. Gregory, Anna C. Haber, Donghee Hoh, Emily E. Jennings, Zhongjie Ji, Prabhjot Kaur, Sunil K. Kenchanmane Raju, Yunfei Long, Serena G. Lotreck, Davis T. Mathieu, Thilanka Ranaweera, Eleanore J. Ritter, Rie Sadohara, Robert Z. Shrote, Kaila E. Smith, Scott J. Teresi, Julian Venegas, Hao Wang, McKena L. Wilson, Alyssa R. Tarrant, Margaret H. Frank, Zoë Migicovsky, Jyothi Kumar, Robert VanBuren, Jason P. Londo, Daniel H. Chitwood

**Affiliations:** Genetics Program, Michigan State University, East Lansing, Michigan 48824 USA; Department of Biochemistry and Molecular Biology, Michigan State University, East Lansing, Michigan 48824 USA; Department of Plant Biology, Michigan State University, East Lansing, Michigan 48824 USA; Department of Horticulture, Michigan State University, East Lansing, Michigan 48824 USA; Department of Plant, Soil and Microbial Sciences, Michigan State University, East Lansing, Michigan 48824 USA; Cell and Molecular Biology Program, Michigan State University, East Lansing, Michigan 48824 USA; MSU-DOE Plant Research Laboratory, Michigan State University, East Lansing, Michigan 48824 USA; Molecular Plant Sciences Program, Michigan State University, East Lansing, Michigan 48824 USA; Plant Breeding, Genetics, and Biotechnology, Michigan State University, East Lansing, Michigan 48824 USA; Department of Electrical and Computer Engineering, Michigan State University, East Lansing, Michigan 48824 USA; Department of Computational Mathematics, Science & Engineering, Michigan State University, East Lansing, Michigan 48824 USA; School of Integrative Plant Science, Plant Biology Section, Cornell University, Ithaca, New York 14850 USA; Department of Plant, Food and Environmental Sciences, Faculty of Agriculture, Dalhousie University, Truro, Nova Scotia B2N 5E3 Canada; Grape Genetics Research Unit, USDA ARS, Geneva, New York 14456 USA

**Keywords:** grapevine, landmark analysis, leaf shape, modeling, morphometrics, Vitis

## Abstract

**Premise of study:** Leaf morphology is dynamic, continuously deforming during leaf expansion and among leaves within a shoot. We measured leaf morphology from over 200 vines over four years, and modeled changes in leaf shape along the shoot to determine if a composite “shape of shapes” can better capture variation and predict species identity compared to individual leaves.

**Methods:** Using homologous universal landmarks found in grapevine leaves, we modeled various morphological features as a polynomial function of leaf node. The resulting functions are used to reconstruct modeled leaf shapes across shoots, generating composite leaves that comprehensively capture the spectrum of possible leaf morphologies.

**Results:** We found that composite leaves are better predictors of species identity than individual leaves from the same plant. We were able to use composite leaves to predict species identity of previously unassigned vines, which were verified with genotyping.

**Discussion:** Observations of individual leaf shape fail to capture the true diversity between species. Composite leaf shape—an assemblage of modeled leaf snapshots across the shoot—is a better representation of the dynamic and essential shapes of leaves, as well as serving as a better predictor of species identity than individual leaves.

## INTRODUCTION

Leaf shape is dynamic. Allometry—the differential growth of shape attributes relative to size or other attributes—was first documented in leaves by Stephen Hales in 1727. Imprinting a grid of points onto expanding young fig leaves led to the observation of “the difference of the progressive and lateral motions of these points in different leaves”, finding that they “were of very different lengths in proportion to their breadths” (Hales, 1727). Not only is the shape of a leaf dynamic over its development as it expands, but that of multiple leaves located at different nodes within a plant is as well. Heteroblasty—phenotypic changes in sequential lateral organs (such as leaves)—was first described by Johann Wolfgang von Goethe (a contributor to art, philosophy, and botany; Friedman and Diggle, 2011) when he compared the transformation of mature leaf shapes within a plant to the Greek god of the sea, and the mutable nature of water (Goethe, 1817; 1952).

Our ability to recognize differences in leaf shape indicates that this is a quantifiable trait. A number of morphometric approaches have been proposed to measure leaf shape across angiosperms, from focusing on the venation using computer vision (Wilf et al., 2016) to the closed contour of the blade using Topological Data Analysis (Li et al., 2018). In some special cases, simple but powerful geometric approaches have been developed to aid in the classification of plants. For example, all grapevine leaves have corresponding, homologous primary and secondary veins. With the introduction of North American rootstocks in Europe— to combat the spread of phylloxera—during the late 19^th^ and early 20^th^ centuries viticulturists were exposed to new and unfamiliar varieties; thus, they needed a method to confirm rootstock identity. Without the ability to genotype (yet), viticulturists used phenotype, and proposed that the angle of the petiolar veins, which form the petiolar sinus, could be used to differentiate between varieties (Goethe 1876; 1878; Ravaz, 1902). This system was extended to other major veins in the leaf, measuring their relative angles and the ratios of lobe and sinus lengths, by Pierre Galet (Galet 1979; 1985; 1988; 1990; 2000). A system of homologous features was seized upon to further elaborate all veins hierarchically and to enumerate the corresponding teeth in which they terminate (Rodrigues 1939; 1941a; 1941b; 1952a; 1952b). María-Carmen Martínez used these approaches to calculate average leaves (Martínez and Grenan, 1999) and with this mathematical framework classified varieties, clones, and even their similarity to depictions of grapevines in art (Martínez et al., 1995; 1997a; 1997b; Santiago et al., 2005; 2007; 2008; Gago et al., 2009a; 2009b; 2014). These morphometric methods, which have been tailored to the unique geometric properties of grapevine leaves for over a century, have been extended to formal landmark-based methods and have been used in the study of the genetic basis of leaf shape (Chitwood et al., 2014; Demmings et al., 2019).

The unique features of grapevine leaves allow not only for the classification of different varieties, but also the study of the dynamics of leaf morphology on individual vines, as these features vary along the grapevine shoot (**Fig. 1**). From shoot base to tip, two developmental processes—allometry and heteroblasty—are discernable through observation of leaf size and shape (**Fig. 2**). Following initiation at the shoot tip, leaves rapidly undergo expansion while their shape continuously deforms, governed by the allometric processes first described by Hales (1727). Heteroblasty is caused by temporal changes in the shoot apical meristem that alter the phenotype of the subsequent lateral organs (in this case, leaves) produced, including leaf shape. Leaves found at the shoot base were the first to emerge from buds; thus, leaves at nodes closer to the base are relatively mature and have reached their maximum size. At this point in development, differences in leaf shape are predominantly attributable to the transformations from “first to last” described by Goethe (1817) between mature leaves with different shapes.

**Figure 1:**
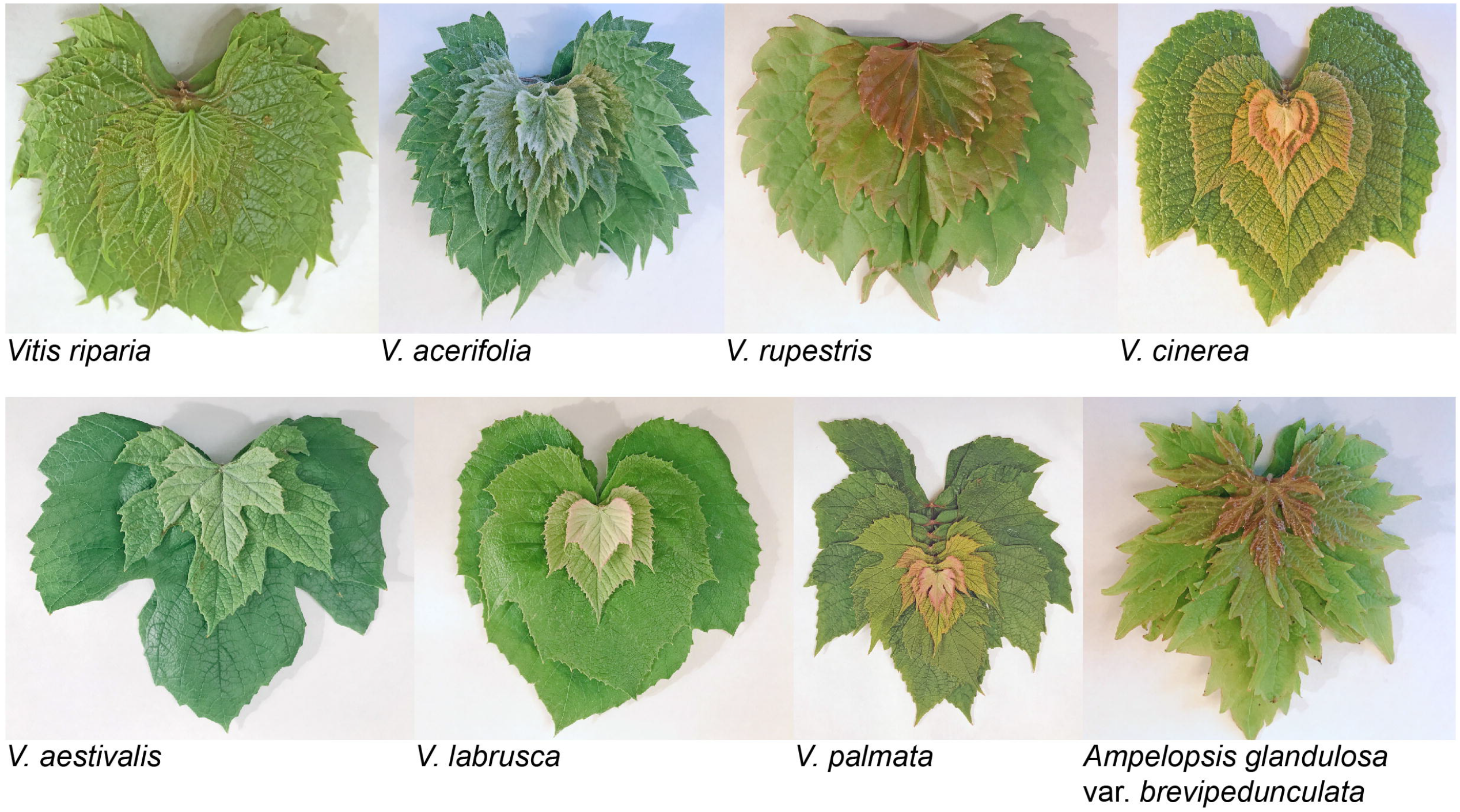
Examples of changes in leaf traits between different developmental stages in different grapevine species. Vineyard-collected leaves (adaxial side up, except for V. riparia) from the tip (top of stack) to base (bottom of stack) of the shoot. Size, shape, and color (among other traits) vary from node to node. Images are not to scale relative to each other.

**Figure 2:**
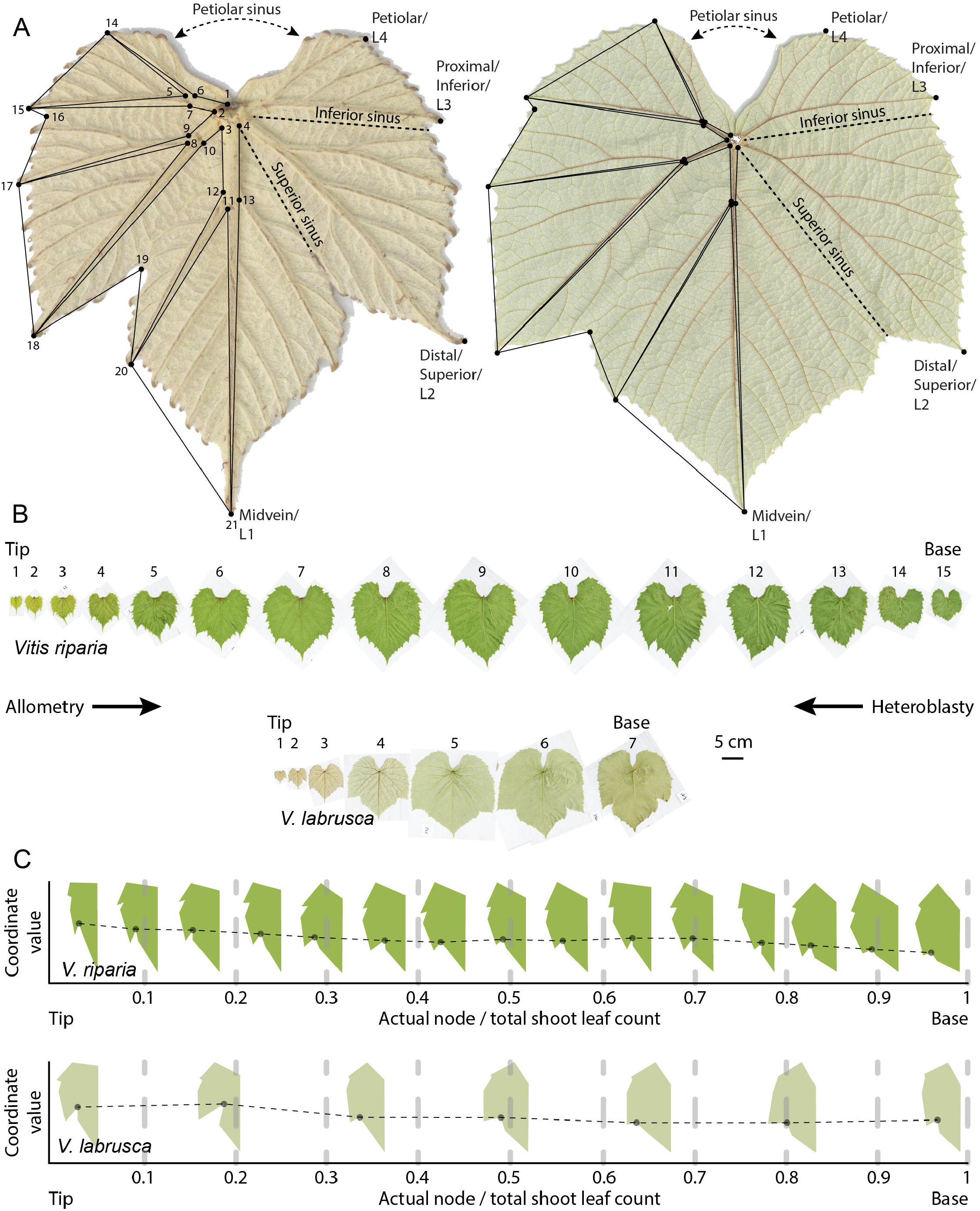
Quantifying leaf shape changes along the shoot. **A)** Two Vitis labrusca leaves from the second (left) and fourth (right) nodes from the shoot tip. Landmarks are indicated to the left, and lobe tips, sinuses, and associated nomenclature to the right. Leaves are scaled to show the allometric decrease in the ratio of vein-to-blade area that occurs during expansion. **B)** Examples of morphological changes in V. riparia and V. labrusca leaves sampled along the shoot. Shoot tip, shoot base, nodes, and scale are indicated. Leaves in (A) are from the V. labrusca shoot shown here. **C)** Diagrammatic representation of the methods used to quantify leaf shape change along the shoot. Leaves are first superimposed and scaled using Procrustean methods. The outlines shown were formed using Procrustean coordinates derived from the blade of each leaf shown in (B). Relative node position is calculated as the node number (starting at the tip) divided by total leaf count (for the shoot), such that all nodes are assigned a fractional value between 0 and 1. Coordinate x- and y-values are modeled as a function of relative node position. Dots connected between leaves by the dotted line correspond to landmark 19 of the distal sinus.

Previously, we measured leaf shape from hundreds of vines from a germplasm collection representative of North American *Vitis* species (Chitwood et al., 2016a). We were able to capture changes in shape between species as well as changes due to allometry and heteroblasty by sampling leaves from all nodes on a single shoot. We found that the effects of species and developmental processes on leaf shape were additive and statistically distinct, meaning that regardless of node chosen along the vine, a given leaf can be used to identify the species and vice versa. We sampled leaves from the same vines in a second season (Chitwood et al., 2016b), and, by comparing equivalent leaves (by the vine and node) between the two growing seasons (2013 and 2015), we observed interannual variability in leaf shape that was attributable to climate rather than genotype or development.

Here, using the same vines as in our previous work (Chitwood et al., 2016a; 2016b), we sampled leaves from two more seasons (2016 and 2017) and successfully use composite leaf-modelling to predict species identity in vines with unknown background, which were confirmed by genotyping. Our work demonstrates that phenotypic modeling of dynamic changes in leaf shape, using composite leaves, improves genotype predictions compared to approaches statistically accounting for individual leaves only.

## METHODS

### Germplasm, sample collection, and imaging

Leaves (8,465 in total) were collected from 209 vines at the USDA germplasm repository vineyard in Geneva, New York (USA). Samples were taken from the same vine during the 2nd week of June, annually, in 2013 and 2015-17. This study builds upon and analyzes previous work, including datasets from 2013 (Chitwood et al., 2016a) and 2013 + 2015 (Chitwood et al., 2016b). The vines sampled represent 11 species (*Ampelopsis glandulosa* var. *brevipedunculata, Vitis acerifolia, V. aestivalis, V. amurensis, V. cinerea, V. coignetiae, V. labrusca, V. palmata, V. riparia, V. rupestris, and V. vulpina), four hybrids (V. ×andersonii, V. ×champinii, V. ×doaniana, and V. ×novae-angliae*), and 13 *Vitis* vines with unassigned identity. Starting at the shoot tip (with shoot order noted for each leaf), leaves greater than ~1 cm in length were collected in stacks (**Fig. 1**) and stored in labelled plastic bags with ventilation holes in a cooler. Within 2 days of collection, leaves were arranged on a large-format Epson Workforce DS-50000 scanner (Tokyo, Japan) in the order collected, with a small number near each leaf indicating which node it came from and a ruler for scale within the image file. Image files are named with the vine ID, followed by a sequential lowercase letter if multiple scans were needed.

### Landmarking and Generalized Procrustes Analysis

Twenty-one landmarks were (manually) annotated sequentially on the abaxial side of the leaf, as in **Fig. 2A**, using the point tool in ImageJ (version 1.52k, Abràmoff et al., 2004). From the 8,465 leaves used in this study, 177,765 landmarks (355,530 values in total) were analyzed. Landmarks were placed sequentially for each leaf in a scan and saved as a text file of *x*- and *y*- coordinate values. To check for errors, landmarks from each scan were visualized using ggplot2 (Wickham, 2016) in R (version 3.6.0, R Core Team, 2019), and landmarking was redone as necessary. Generalized Procrustes Analysis (GPA) was performed using the shapes package (version 1.2.4, Dryden and Mardia, 2016) in R (R Core Team, 2019), allowing for reflection. The resulting superimposed Procrustes coordinates were used in subsequent analyses.

### Data analyses

The data analysis methods used in this study were devised by students during Fall 2019 in *Foundation in Computational Plant Science*, a graduate-level course offered through the Department of Horticulture at Michigan State University, which focused on the integration of computational and plant science approaches. Use of the “flipped classroom” approach, Jupyter notebooks (Kluyver et al., 2016), and the Python coding language to teach statistical and modeling analyses was inspired by *Introduction to Computational Modeling and Data Analysis*, an undergraduate course offered through the Department of Computational Mathematics, Science & Engineering at Michigan State University (Silvia et al., 2019; 2020). The learning objectives of both courses focus on teaching coding (assuming no prior experience) in addition to principles of statistics and modeling.

An innovative feature of Jupyter notebooks is their multifunctional use in both education and research. Additionally, as an open-source platform, Jupyter notebooks facilitate sharing between multiple sources, including the classroom and laboratory. Creativity and skill drive data-analysis oriented research; thus, educational methods are essential and should be shared, reproduced, and improved upon.

All Python code (Jupyter notebooks) and R scripts used for data analysis are available on github: https://github.com/DanChitwood/grapevine_shoots. The concept of modeling Procrustean coordinate values across nodes using polynomial functions was introduced to students using published leaf shape data from Passiflora spp. (Chitwood and Otoni, 2017a; 2017b): https://github.com/DanChitwood/PlantsAndPython/blob/master/PlantsAndPython10_STUDENT_A_Passion_for_Passiflora.ipynb. Jupyter notebooks used as instructional materials for the class are available in the *PlantsAndPython* repository: https://github.com/DanChitwood/PlantsAndPython.

Analyses and visualizations in Python were done with NumPy (Oliphant, 2006), pandas (McKinney, 2010), scikit-learn (Pedregosa et al., 2011), and Matplotlib (Hunter, 2007). In order to compare node position of each leaf against each other, we created a relative node position, which is simply the node position for each leaf divided by the total leaf count for the shoot, where 0 is the shoot tip and 1 is the shoot base (**Fig. 2**). Procrustean coordinates for each vine were modeled as a function of relative node using a second-degree polynomial function fitted to the data using the NumPy polyfit and poly1d functions (**Appendix S1**). Ten modeled leaf shapes were calculated across the normalized range of node values from zero to one for each shoot. Collectively, the coordinate values for these ten modeled leaf shapes were used in subsequent analyses, representing leaf shape changes across the shoot as a composite leaf shape. Principal Component Analysis (PCA) and Linear Discriminant Analysis (LDA) were performed using scikit-learn. For LDA and classification, random resampling was used to even the replication between species. For the test set, 20% of the data was used, and the remaining 80% was used for training. The genotypes of *Vitis* spp. and parentage of *Vitis* hybrids were unknown (or subject to speculation); thus, these individuals were omitted from the training set but included in the test set, to determine which species their leaves most resembled. LDA prediction was run 1,000 times to estimate precision, recall, accuracy, and F1 statistics.

### Genotypic data and ADMIXTURE analysis

Genotype data is derived from Klein et al. (2018). VCF files containing genotype data for the grapevines were processed using PLINK2 (Chang et al., 2015), resulting in a binary biallelic genotype table (.bed), PLINK extended MAP file (.bim), and PLINK sample information file (.fam). PLINK2 was used to calculate eigenvectors and eigenvalues used in PCA (Galinsky et al., 2016). The biallelic genotype table was used in a single run of ADMIXTURE (Alexander et al., 2009). K-values from 3-15 were run and CV error was calculated. CV error decreased consistently from 0.20442 with K = 3 to 0.16453 with K = 10, after which it fluctuated around 0.164 for K values of 11-15. For final analysis, K = 10 was used. The resulting table of group proportions was analyzed in R and visualized with ggplot2.

## RESULTS

### Using composite leaves to model leaf shape along the shoot

Vein area relative to that of the whole leaf blade decreases exponentially with leaf expansion in grapevine, making this trait an important allometric feature of leaf morphology (**Fig. 2A**; Chitwood et al., 2016b). We measured leaf shape and vein width using 21 homologous landmarks, which were superimposed using a Generalized Procrustes Analysis such that the resulting coordinates for each sample were translated, rotated, reflected, and scaled; thus, allowing for cross-comparison of shape (Gower, 1975). Relative node position was used to compare vines with different numbers of leaves (**Fig. 2B-C**).

With comparable Procrustes-adjusted coordinates and relative node positions, the coordinate *x*- and *y*-values for each of the 21 homologous landmarks (42 per leaf) were modeled as a function of node position. There was never more than one inflection point and the data were smooth; thus, a second order polynomial function was used for modelling—also selected so as to not overfit the data, which can occur with higher-order polynomials (**Appendix S1**). Using the resulting functions, coordinate values for 10 modeled leaf shapes were calculated for each vine, in intervals of 0.1 across the relative node space from 0.1-1. The resulting modeled leaf shapes (with 210 landmarks and 420 coordinate values) are referred to as composite leaves and represent the dynamic changes in shape across each shoot sampled.

Composite leaves can be superimposed and visualized, comparing changes in leaf shape along the shoot in different species, effectively reflecting both genotypic (species) and developmental (node position) differences (**Fig. 3**). Some of the most noticeable differences in leaf shape were observed between different species—the wide reniform leaves of *Vitis rupestris*, the broad orbicular leaves of *V. labrusca, V. coignetiae, and V. amurensis*, and the deep-lobed leaves of *V. palmata* and *Ampelopsis glandulosa var. brevipedunculata*. Developmental trends were also observed. Generally, the youngest leaves (at the shoot tip) were thinner and had deeper lobes than mature leaves found at the shoot base. Composite leaves were able to capture the dynamic developmental changes in leaf shape along the shoot, which individual leaves are not able to do, a “shape of shapes” that better represents leaf shape across different species.

**Figure 3:**
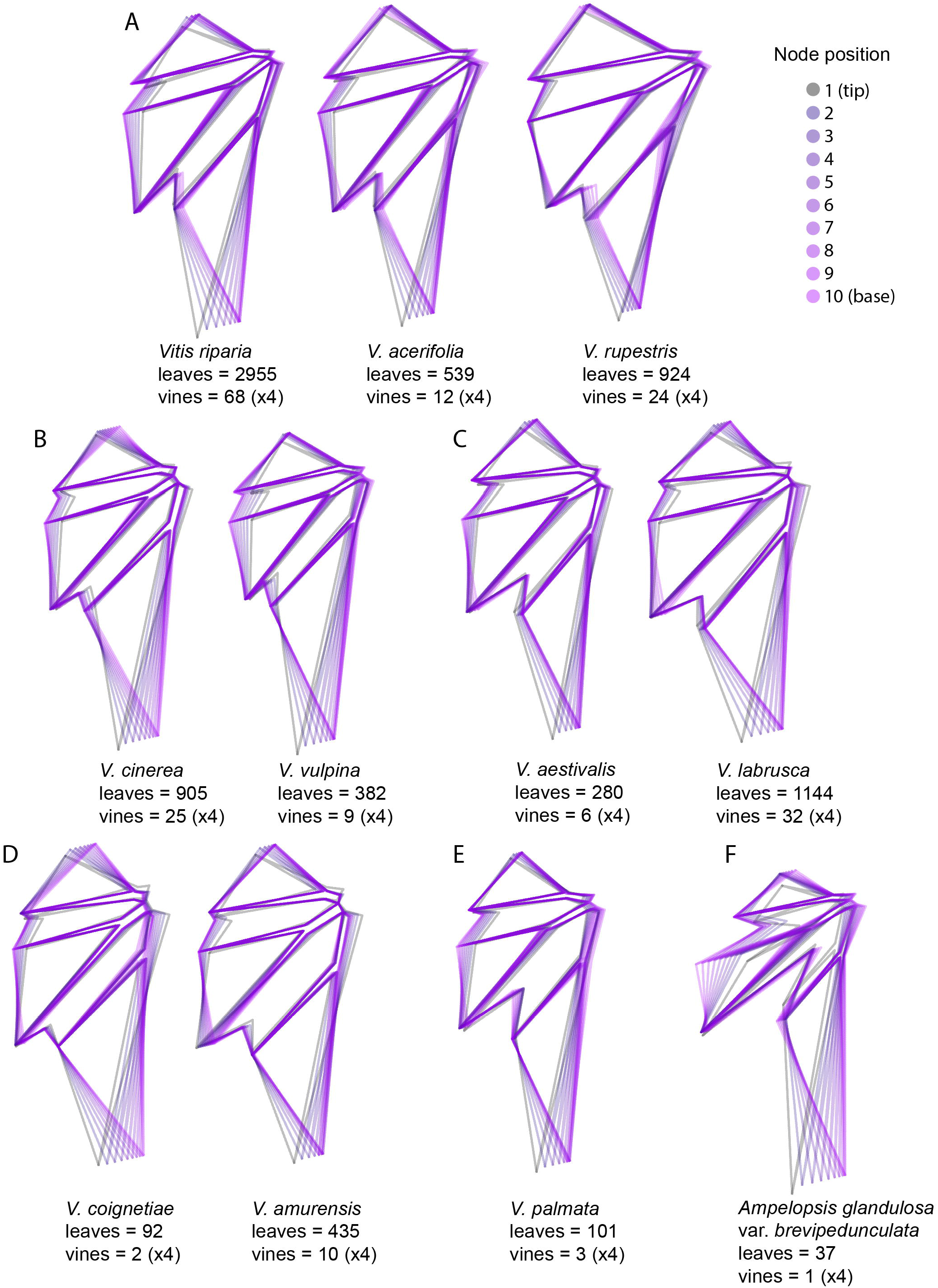
Modeling leaf shape along grapevine shoots with composite leaves. For each species, modeled leaf shapes for ten relative node positions along the shoot were superimposed and illustrated, forming a composite leaf. Illustrations are grouped by species relatedness: **A)** Vitis riparia, V. acerifolia, V. rupestris; **B)** V. cinerea, V. vulpina; **C)** V. aestivalis, V. labrusca; **D)** V. coignetiae, V. amurensis; **E)** V. palmata; **F)** Ampelopsis glandulosa var. brevipedunculata. For each species, the number of leaves and vines sampled is given (note that every vine is sampled across four years, yielding a pseudoreplication of four). Composite leaves are colored as a gradient from gray (the shoot tip, node 1) to pink-purple (the shoot base, node 10).

### Composite leaves outperform individual leaves in predicting species identity

We hypothesized that composite leaves would outperform individual leaves in predicting species identity. To test this, we performed a Principal Component Analysis (PCA) on individual vs. composite leaf shape. Performing PCA for individual leaves permits representation of the axes of variation along Principal Components (PCs) as eigenleaves (theoretical leaf shapes representing variation along each PC axis at a chosen standard deviation value; **Fig. 4A**). A set of 4 example species with distinct leaf morphologies (*Vitis riparia, V. acerifolia, V. rupestris*, and *Ampelopsis glandulosa var. brevipedunculata*) was compared, and the individual-leaf PCA separated species predominantly by PC2 (**Fig. 4B**). The eiglenleaf representations support this structure, as a short and wide reniform leaf type, characteristic of *V. rupestris*, is associated with low PC2 values, whereas the longer leaf type with more prominent lobes more similar to*A. glandulosa* var. *brevipedunculata* is associated with high PC2 values. In contrast, the separation between species is much greater for the composite leaf PCA space (**Fig. 4C**).

**Figure 4:**
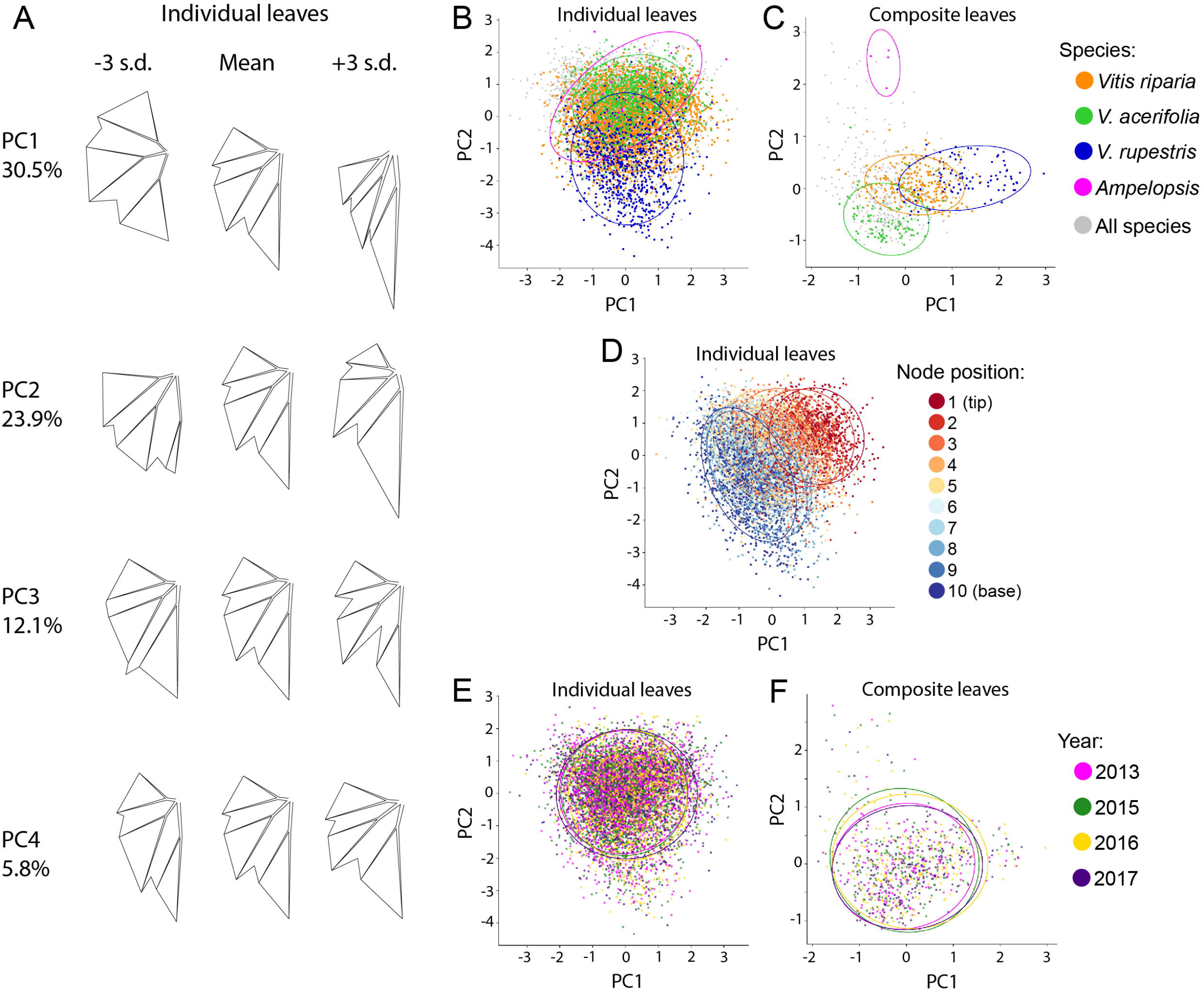
Principal Component Analysis (PCA) of individual vs. composite leaves. **A)** Eigenleaf representations of shape variance at −/+ 3 standard deviations explained by Principal Components (PCs) for a PCA performed on all individual leaves. The percent variance explained by each PC is shown (on the left). **B-C)** Comparison of results from two separate PCAs, one with all individual (B) and the other with composite (C) leaves. Confidence ellipses (95%) for four species (following the color legend) are provided in addition to all data points (gray). **D)** Relative node position discretized into nodes counting from one to ten projected onto the individual leaf PCA space. For relative node position, there is no composite leaf PCA as nodes are accounted for and integrated into the resulting values for that analysis. **E-F)** Individual (E) and composite (F) leaf PCAs with 95% confidence ellipses for each year (following the color legend).

Node position (discretized into ten relative nodes) could only be projected for individual leaf PCA, as it is intrinsic to the modeling approach used. Node position varied mostly by PC1 in individual leaf PCA (**Fig. 4D**) associated with eigenleaf representations characteristic of the shoot tip (high PC1 values) and the shoot base (low PC1 values). The species factor varied along one axis (PC2) and node position along another (PC1), consistent with previous observations that species and developmental effects on leaf shape are additive and orthogonal (Chitwood et al., 2016a). The main morphological difference between the two axes is the association of the petiolar sinus with other features (**Fig. 4A**). Four replicates for each vine are included in the composite leaf space, corresponding to the four growing seasons during which each vine was sampled. Comparing individual (**Fig. 4E**) and composite (**Fig. 4F**) leaf PCA with data separated by year yielded little separation, as is expected given that the main source of variance (species type and developmental effects) was balanced in each analysis.

The increased separation of species in composite compared to individual leaf PCA (**Fig. 4B-C**) suggests that composite leaves may discriminate between species better than individual leaves. To test the ability of these two methods to predict species identity, we used Linear Discriminant Analysis (LDA). The resulting confusion matrices, which plot the proportion of predicted species (horizontal axis) for each actual species class (vertical axis), showed that composite leaves outperform individual leaves in predicting species identity (**Fig. 5A-B**). Precision (true positives divided by the total number of positive predictions), recall (true positives divided by true positives + false negatives), and the F1 score (the harmonic mean of precision and recall) for species prediction were higher for composite compared to individual leaves (**Table 1**). For species prediction, the minimum values of precision, recall, accuracy, and the F1 score were 0.57, 0.50, 0.50, and 0.53, respectively, for individual leaves, and 0.85, 0.84, 0.84, and 0.86 for composite leaves, demonstrating the superior predictive ability of composite leaves. Node position can be predicted for individual leaves but not for composite leaves. For individual leaves, prediction was the most accurate at the shoot tip and base (**Fig. 5C**; **Table 2**), as the effects of allometry and heteroblasty are most pronounced at the tip and base of the shoot, respectively, with little influence over leaves in the middle. Although matched by vine and species, the prediction of year also showed improvement with composite compared to individual leaves (**Fig. 5D-E**; **Table 3**). This indicates that composite leaves retain morphological information useful for discriminating effects more subtle than genotype or development, such as the environment.

**Figure 5:**
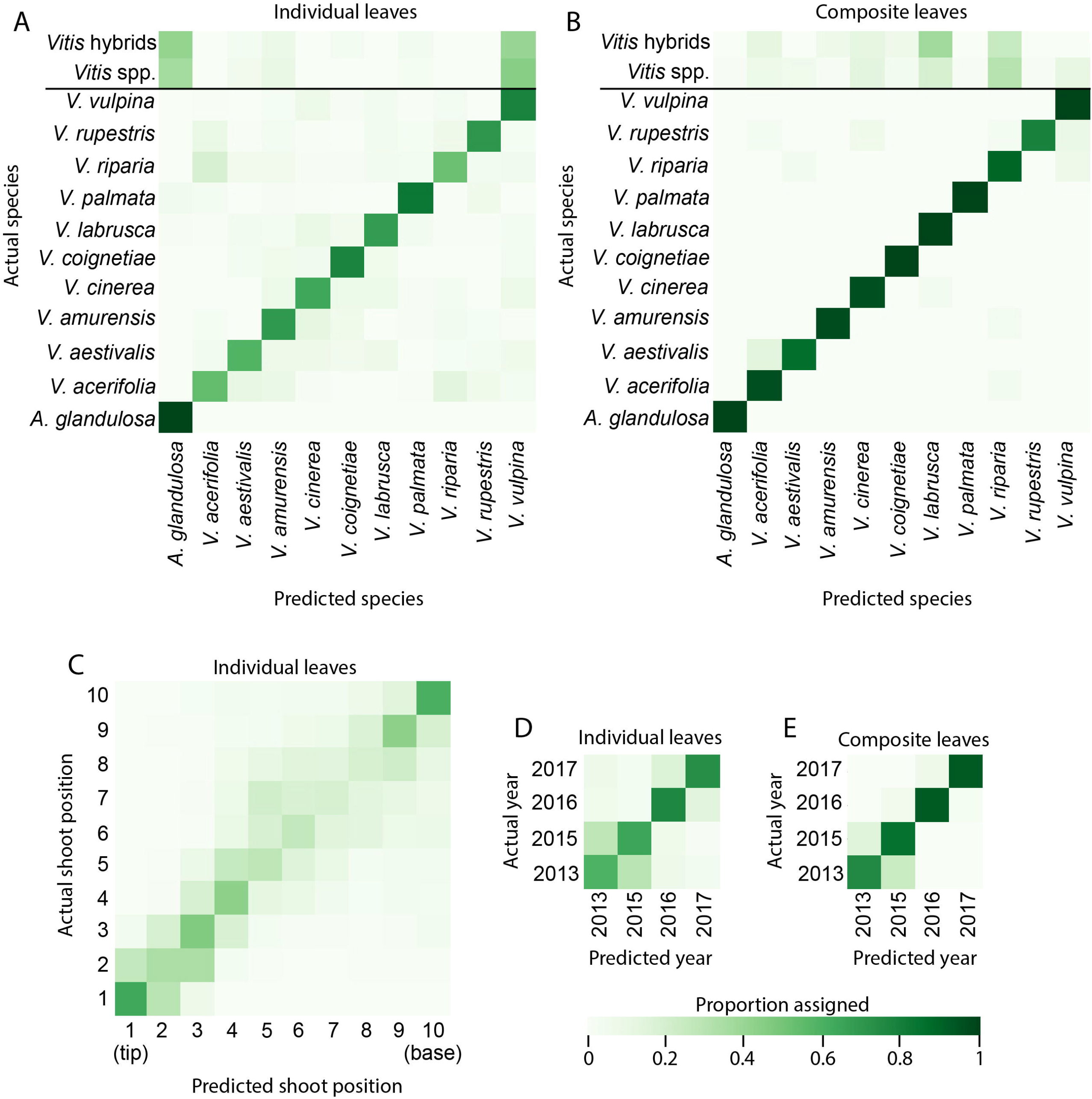
Comparison of Linear Discriminant Analysis (LDA) results for individual vs. composite leaves. **A-B)** Comparison of confusion matrices from two separate LDAs, one for individual (A) and the other for composite (B) leaves. The proportion of actual species (vertical) assigned to predicted species identity (horizontal) is indicated by color. Vitis hybrids and species were not used in the training set and were only assigned identity in the test set. **C)** Confusion matrix for an LDA performed on relative node position discretized into ten nodes along the shoot for individual leaves. For relative node position, there is no composite leaf LDA as nodes are accounted for and integrated into the resulting values for that analysis. **D-E)** Individual (D) and composite (E) leaf LDAs predicting year. All panels use the indicated color scheme for assigned proportion from 0 (white) to 1 (dark green).

**Table 1:** Comparison of species prediction using LDA for individual vs. composite leaves.

| Species | Individual leaves |  |  |  | Composite leaves |  |  |  |
| --- | --- | --- | --- | --- | --- | --- | --- | --- |
|  | Precision | Recall | Accuracy | F1 | Precision | Recall | Accuracy | F1 |
| *A. glandulosa* | 0.93 | 1.00 | 1.00 | 0.96 | 1.00 | 1.00 | 1.00 | 1.00 |
| *V. acerifolia* | 0.57 | 0.50 | 0.50 | 0.53 | 0.90 | 0.94 | 0.94 | 0.92 |
| *V. aestivalis* | 0.68 | 0.56 | 0.56 | 0.62 | 0.96 | 0.97 | 0.97 | 0.97 |
| *V. amurensis* | 0.65 | 0.73 | 0.73 | 0.69 | 0.99 | 0.97 | 0.97 | 0.98 |
| *V. cinerea* | 0.59 | 0.69 | 0.69 | 0.64 | 0.97 | 0.91 | 0.91 | 0.94 |
| *V. coignetiae* | 0.74 | 0.77 | 0.77 | 0.75 | 1.00 | 1.00 | 1.00 | 1.00 |
| *V. labrusca* | 0.67 | 0.65 | 0.65 | 0.66 | 0.95 | 0.97 | 0.97 | 0.96 |
| *V. palmata* | 0.85 | 0.84 | 0.84 | 0.85 | 1.00 | 1.00 | 1.00 | 1.00 |
| *V. riparia* | 0.58 | 0.51 | 0.51 | 0.54 | 0.85 | 0.87 | 0.87 | 0.86 |
| *V. rupestris* | 0.76 | 0.73 | 0.73 | 0.75 | 0.99 | 0.84 | 0.84 | 0.91 |
| *V. vulpina* | 0.69 | 0.75 | 0.75 | 0.72 | 0.88 | 0.99 | 0.99 | 0.93 |

**Table 2:** Relative node position prediction using LDA for individual leaves.

| Node position | Precision | Recall | Accuracy | F1 |
| --- | --- | --- | --- | --- |
| 1 (tip) | 0.66 | 0.66 | 0.66 | 0.66 |
| 2 | 0.42 | 0.37 | 0.37 | 0.39 |
| 3 | 0.41 | 0.44 | 0.44 | 0.42 |
| 4 | 0.35 | 0.40 | 0.40 | 0.37 |
| 5 | 0.27 | 0.30 | 0.30 | 0.28 |
| 6 | 0.26 | 0.26 | 0.26 | 0.26 |
| 7 | 0.29 | 0.22 | 0.22 | 0.25 |
| 8 | 0.28 | 0.23 | 0.23 | 0.25 |
| 9 | 0.35 | 0.39 | 0.39 | 0.36 |
| 10 (base) | 0.53 | 0.56 | 0.56 | 0.54 |

**Table 3:** Comparison of year prediction using LDA for individual vs. composite leaves.

| Year | Individual leaves |  |  |  | Composite leaves |  |  |  |
| --- | --- | --- | --- | --- | --- | --- | --- | --- |
|  | Precision | Recall | Accuracy | F1 | Precision | Recall | Accuracy | F1 |
| 2013 | 0.58 | 0.56 | 0.56 | 0.57 | 0.83 | 0.79 | 0.79 | 0.80 |
| 2015 | 0.62 | 0.64 | 0.64 | 0.63 | 0.79 | 0.85 | 0.85 | 0.82 |
| 2016 | 0.74 | 0.79 | 0.79 | 0.76 | 0.95 | 0.96 | 0.96 | 0.95 |
| 2017 | 0.79 | 0.73 | 0.73 | 0.76 | 0.97 | 0.95 | 0.95 | 0.96 |

### Using composite leaves to predict genotype and phenotype

We demonstrated that composite leaves outperform individual leaves in discriminating vines of known species identity. We subsequently sought to test the predictive ability of composite leaves on vines of unassigned identity. Of the 209 vines measured, 147 had been genotyped previously (Klein et al., 2018). Vitis vines of unassigned species identity totaled 13, 10 of which have been genotyped. ADMIXTURE analysis of genotyped vines confirmed most known species groups (Moore, 1991; Miller et al., 2013), with some exceptions (**Fig. 6**). Some intraspecies population structure was detected in *V. riparia* and *V. acerifolia* vines. *V. amurensis* and *V. coignetiae* had low replication and are not resolved (Fig. 6A). *V. aestivalis* vines were either misidentified from *V. palmata* or are unresolved hybrids of *V. palmata* × (*V. labrusca* + *V. aestivalis*). A number of other misassigned vines are indicated by small roman numerals in **Fig. 6A**. These and *V. aestivalis* vines occupy positions in the PCA genotype space inconsistent with their assigned identities or between species groups reflecting complex ancestry (**Fig. 6C-D**).

**Figure 6:**
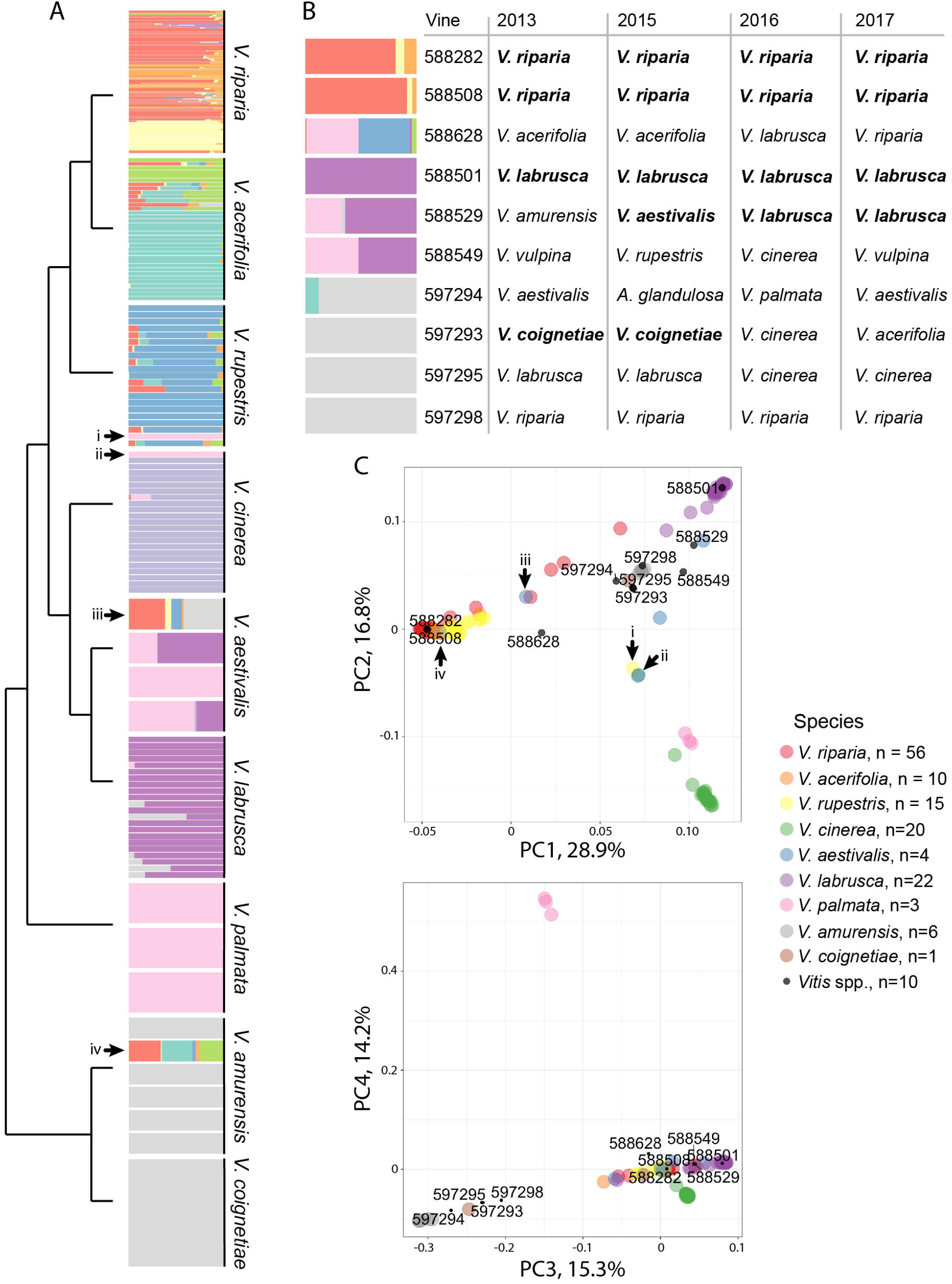
Comparing species identity predictions based on morphology to known ancestry. **A)** Ancestry for each individual using K = 10 from ADMIXTURE. Each population is assigned a different color. Species designations for each vine are as previously assigned for this collection, without prior genetic knowledge, and arranged by known phylogenetic relationships. Vines with genetic identities at odds with their assigned identity are indicated by black arrows and lowercase roman numerals. **B)** For Vitis spp. with genetic information, ancestry (left) and predicted species identity (based on morphology) for each shoot for each of four years (right) is provided. Morphological predictions consistent with genetic identity are indicated in bold. **C)** Principal Component Analysis (PCA) of the same individuals in (A). Vines with conflicting assigned and genetic identities are indicated by black arrows and lowercase roman numerals as in (A). Vitis spp. in (B) are indicated by black dots and vine identification numbers. Species are indicated by colors which do not correspond with the color scheme of other panels and the number of vines with genetic information is provided.

For each *Vitis* spp. vine with genotype data, we predicted species identity using composite leaves with our LDA classifier (**Fig. 5**) for each of the four years sampled (**Fig. 6B**). Three vines were unambiguously identified using leaf shape consistently across the four years: vines 588282 and 588508 were correctly predicted to be *V. riparia*, and vine 588501 as *V. labrusca*.

These vines clustered clearly with their respective species groups within the genotype PCA space (**Fig. 6C-D**). Two additional vines were ambiguously identified: vine 588529 has a (*V. labrusca* + *V. aestivalis*) × *V. palmata* genotype characteristic of *V. aestivalis* and was identified as *V. labrusca* or *V. aestivalis* in three of the four years. Vine 597293 with a (*V. amurensis* + *V. coignetiae*) ancestry was identified as *V. coignetiae* in two of the four years. Vine 588549 was not predicted correctly using composite leaf shape and has a (*V. labrusca* + *V. aestivalis*) × *V. palmata* genotype similar to vine 588529, but lies even farther from the *V. labrusca* cluster in genotype PCA space, suggesting that the more ambiguous the genotype of a vine from a well characterized species, the more difficult morphological-based prediction is.

Vine 588628 had a unique *V. rupestris* × *V. palmata* ancestry and occupied an ambiguous position in the genotypic PCA. We expect that this vine was not classified correctly using morphology. The remaining vines that were not predicted are all of (*V. amurensis* + *V. coignetiae*) ancestry (vines 597294, 597295, and 597298). Likely, because of the small sample sizes for these two species, we have not adequately sampled leaf shape for this lineage, and therefore, cannot accurately predict for test cases. Our results show that composite leaf shape can predict species identity successfully, but not for vines with complex ancestry or species that have not been adequately sampled for leaf shape variation.

## DISCUSSION

Previously, we analyzed leaf morphology across nodes in different species of *Vitis* and *Passiflora* and found that allometric and heteroblastic effects are statistically separable from species specific differences (Chitwood et al., 2016a; Chitwood and Otoni, 2017a). The tracking of alterations to leaf shape between nodes was critical for several observations, including 1) the determination that species-specific differences in leaf shape arise from a common juvenile form (Chitwood and Otoni, 2017b) and 2) the identification of alterations to leaf morphology between growing seasons/years (Chitwood et al., 2016b; Baumgartner et al, 2020). However, in these instances, individual leaf shape was statistically analyzed, meaning that node position is a statistical effect rather than being part of the phenomenon studied (shape). By normalizing nodes against overall leaf count per shoot (**Fig. 2B-C**) and subsequently modeling Procrustes adjusted coordinates as a function of node position (Appendix S1), we are able to construct composite leaves—a shape of shapes (**Fig. 3**), able to capture multiple leaf-forms along the shoot in a single object.

Shifting the concept of leaf shape from singular leaves to multiple, sequential, leaves found along the shoot has repercussions for the morphological species concept and phenotype. Species are better-resolved and predicted morphologically with composite (**Fig. 4**) rather than individual leaves alone (**Fig. 5**). With an adequate training dataset, composite leaves can help to identify vines with unknown species identity (**Fig. 6**). Leaf shape—a feature that is complex, conspicuous, and easily observable—is readily used to classify closely-related botanical specimens, as well as assess developmental- or environmentally-driven alterations within single plants. While the diversity of leaf shape can be recognized by specialists and communicated to others through writing, the geometric properties of leaf shape allow for quantification of differences that we perceive visually (Amézquita et al., 2020). Furthermore, our assessment of leaf morphology is limited with individual leaves, which allow us to observe only facets of the comprehensive phenotype. Thus, composite leaves can better help identify and define species by allowing us to capture dynamic morphological data from developmental and environmental conditions compared to individual leaves.

## ACKNOWLEDGEMENTS

DHC and JPL were recipients of a Grape Research Coordination Network stipend from the National Science Foundation, which funded this research along with the USDA National Institute of Food and Agriculture and Michigan State University AgBioResearch. DHC, ZM, and JPL were supported by the National Science Foundation Plant Genome Research Program, award number 1546869. The class in which students that worked on this project participated was supported by the National Science Foundation Research Traineeship Program (NRT), award number 1828149.

## AUTHOR CONTRIBUTIONS

DHC and JPL conceived of the project and experimental design. DHC, JM, ZM, and MF collected data. DHC, RV, and JK developed the curriculum and innovated the teaching methods used to guide student-contributions to the manuscript. All authors contributed to data analysis. DHC wrote the first draft of the manuscript, which was read and edited by all authors.

## DATA AVAILABILITY

All data and code used in this study can be found on github (https://github.com/DanChitwood/grapevine_shoots).

## SUPPORTING INFORMATION

## Appendix S1

Modeled coordinate values across grapevine shoots.

## LITERATURE CITED

Abràmoff, M. D., P. J. Magalhães, S. J. Ram. 2004. Image processing with ImageJ. Biophotonics International 11: 36–42.

Alexander, D. H., J. Novembre, K. Lange. 2009. Fast model-based estimation of ancestry in unrelated individuals. Genome Research 19: 1655–1664.

Amézquita, E. J., M. Y. Quigley, T. Ophelders, E. Munch, D. H. Chitwood. 2020. The shape of things to come: Topological data analysis and biology, from molecules to organisms. Developmental Dynamics.

Baumgartner, A., M. Donahoo, D. H. Chitwood, D. J. Peppe. 2020. The influences of environmental change and development on leaf shape in Vitis. American Journal of Botany 107: 1–13.

Chang, C. C., C. C. Chow, L. C. Tellier, S. Vattikuti, S. M. Purcell, J. J. Lee. 2015. Second-generation PLINK: rising to the challenge of larger and richer datasets. Gigascience 4: s13742–015.

Chitwood, D. H, A. Ranjan, C. C. Martinez, L. R. Headland, T. Thiem, R. Kumar, M. F. Covington, T. Hatcher, D. T. Naylor, S. Zimmerman et al. 2014. A modern ampelography: a genetic basis for leaf shape and venation patterning in grape. Plant Physiology 164: 259–272.

Chitwood, D. H., L. L. Klein, R. O’Hanlon, S. Chacko, M. Greg, C. Kitchen, A. J. Miller, J. P. Londo. 2016a. Latent developmental and evolutionary shapes embedded within the grapevine leaf. New Phytologist 210: 343–355.

Chitwood, D. H., S. M. Rundell, D. Y. Li, Q. L. Woodford, T. Y. Tommy, J. R. Lopez, D. Greenblatt, J. Kang, J. P. Londo. 2016b. Climate and developmental plasticity: interannual variability in grapevine leaf morphology. Plant Physiology 170: 1480–1491.

Chitwood, D. H., W. C. Otoni. 2017a. Morphometric analysis of Passiflora leaves: the relationship between landmarks of the vasculature and elliptical Fourier descriptors of the blade. GigaScience 6: giw008.

Chitwood, D. H., W. C. Otoni. 2017b. Divergent leaf shapes among Passiflora species arise from a shared juvenile morphology. Plant Direct 1: e00028.

Demmings, E. M., B. R. Williams, C. R. Lee, P. Barba, S. Yang, C. F. Hwang, B. I. Resich, D. H. Chitwood, J. P. Londo. 2019. Quantitative Trait Locus Analysis of Leaf Morphology Indicates Conserved Shape Loci in Grapevine. Frontiers in Plant Science 10: 1373.

Dryden, I. L., K. V. Mardia. 2016. Statistical shape analysis: with applications in R (Vol. 995). Hoboken, New Jersey, USA: John Wiley & Sons.

Friedman, W. E., P. K. Diggle. 2011. Charles Darwin and the origins of plant evolutionary developmental biology. The Plant Cell 23: 1194–1207.

Gago, P., J. L. Santiago, S. Boso, V. Alonso-Villaverde, M. S. Grando, M. C. Martínez. 2009a. Biodiversity and characterization of twenty-two Vitis vinifera L. cultivars in the Northwestern Iberian Peninsula. American journal of enology and viticulture 60: 293–301.

Gago, P., J. L. Santiago, S. Boso, V. Alonso-Villaverde, M. C. Martinez. 2009b. Grapevine (Vitis vinifera L.): Old varieties are reflected in works of art. Economic Botany 63: 67–77.

Gago, P., S. Boso, V. Alonso-Villaverde, J. L. Santiago, M. C. Martínez. 2014. Works of Art and Crop History: Grapevine Varieties and the Baroque Altarpieces. Economic Botany 68: 153–168.

Galet, P. 1979. A practical ampelography (L.T. Morton, Trans.). Ithaca, USA: Cornell University Press.

Galet, P. 1985. Précis d’ampélographie pratique, 5 ed., Montpellier, France: Déhan.

Galet, P. 1988. Cépages et vignobles de France, vol. I, Les vignes américaines. Montpellier, France: Déhan.

Galet, P. 1990. Cépages et vignobles de France, vol. II. L’ampélographie française. Montpellier, France: Déhan.

Galet, P. 2000. Dictionnaire encyclopédique des cépages. Paris, France: Hachette.

Galinsky, K. J., G. Bhatia, P. R. Loh, S. Georgiev, S. Mukherjee, N. J. Patterson, A. L. Price. 2016. Fast principal-component analysis reveals convergent evolution of ADH1B in Europe and East Asia. The American Journal of Human Genetics 98: 456–472.

Goethe, J. W. 1817. Goethe’s Werk, Italienische Reise. Stuttgart, Germany: Dreizehnter Band.

Goethe, H. 1876. Note sur l’ampelographie. Congress of Marburg, September 18w.

Goethe, H. 1878. Handbuch der Ampelographie. Graz, Austria: Commission-Verlag von Leykam-Josefsthal.

Goethe J. W. 1952. Botanical Writings, translated by Bertha Mueller, with an introduction by Charles J. Engard. Honolulu, HI: University of Hawaii Press.

Gower, J. C. 1975. Generalized Procrustes Analysis. Psychometrika 40: 33–51.

Hales, S. 1727. Vegetable Staticks: Or, an Account of Some Statical Experiments on the Sap in Vegetables: Being an Essay Towards a Natural History of Vegetation. Also, a Specimen of an Attempt to Analyse the Air, by a Great Variety of Chymio-statical Experiments; which Were Read at Several Meetings Before the Royal Society (Vol. 1). p. 344, Figs. 43-45. London, England: W. and J. Innys and T. Woodward.

Hunter, J. D. 2007. Matplotlib: A 2D graphics environment. Computing in Science & Engineering 9: 90–95.

Klein, L. L., A. J. Miller, C. Ciotir, K. Hyma, S. Uribe-Convers, J. Londo. 2018. High-throughput sequencing data clarify evolutionary relati onships among North American Vitis species and improve identification in USDA Vitis germplasm collections. American Journal of Botany 105(2): pp. 215–226.

Kluyver, T., B. Ragan-Kelley, F. Pérez, B. E. Granger, M. Bussonnier, J. Frederic, K. Kelley, J. B. Hamrick, J. Grout, S. Corlay, P. Ivanov. 2016. Jupyter Notebooks-a publishing format for reproducible computational workflows. In ELPUB. pp. 87–90.

Li, M., H. An, R. Angelovici, C. Bagaza, A. Batushansky, L. Clark, V. Coneva, M. J. Donoghue, E. Edwards, D. Fajardo, et al. 2018. Topological data analysis as a morphometric method: using persistent homology to demarcate a leaf morphospace. Frontiers in Plant Science 9: 553.

Martínez, M. C., M. D. Loureiro, J. L.G. Mantilla. 1995. Importancia y validez de distintos parámetros ampelométricos de hoja adulta para la diferenciación de cepas de Vitis vinifera L., de distintos cultivares. Invest. Agr. Prod. Prot. Veg. 9: 377–389.

Martínez, M. C., J. M. Boursiquot, S. Grenan, R. Boidron. 1997a. *Étude ampelométrique de feuilles adultes de somaclones du cv. Grenache* N (Vitis vinifera L.). Can. J. Bot. 75: 333–345.

Martínez, M. C., S. Grenan, J. M. Boursiquot. 1997b. Variabilidad de algunos caracteres ampelográficos y de producción, en somaclones del cultivar Grenache N (Vitis vinifera L.). Acta Hortic. 18: 271–280.

Martinez, M. S., S. Grenan. 1999. A graphic reconstruction method of an average vine leaf. Agronomie, EDP Sciences 19: 491–507.

McKinney, W. 2010. Data structures for statistical computing in python. In Proceedings of the 9th Python in Science Conference. Vol. 445, pp. 51–56.

Miller, A. J., N. Matasci, H. Schwaninger, M. K. Aradhya, B. Prins, G. Y. Zhong, C. Simon, E. S. Buckler, S. Myles. 2013. Vitis phylogenomics: hybridization intensities from a SNP array outperform genotype calls. PLOS One 8: e78680.

Moore, M. O. 1991. Classification and systematics of eastern North American Vitis L. (Vitaceae) north of Mexico. Sida, contributions to botany, 339–367.

Oliphant, T. E. 2006. A guide to NumPy (Vol. 1, p. 85). USA: Trelgol Publishing.

Pedregosa, F., G. Varoquaux, A. Gramfort, V. Michel, B. Thirion, O. Grisel, M. Blondel, P. Prettenhofer, R. Weiss, V. Dubourg, J. Vanderplas. 2011. Scikit-learn: Machine learning in Python. Journal of Machine Learning Research 12: 2825–2830.

R Core Team. 2019. R: A language and environment for statistical computing. R Foundation for Statistical Computing, Vienna, Austria. Website: https://www.R-project.org/ [accessed 16 April 2020]

Ravaz, L. 1902. *Les vignes américaines: Porte-greffes et producteurs directs*. Montpellier and Paris, France: Goulet. Digitized by Google Books from Cornell University. Public domain. pp. 14, 17–18, 20.

Rodrigues, A. 1939. Sôbre a caracterizaçao das espécies e hibridos do género Vitis. Agron. Lusitana 1: 315–326.

Rodrigues, A. 1941a. Variaçoes do recorte da fôlha da videira. Agron. Lusitana 3: 189–193.

Rodrigues, A. 1941b. Acêrca do valor taxonomico do numero de dentes da fôlha na separaçao de dois hibridos do genero Vitis. L. Agron. Lusitana 3: 325–340.

Rodrigues, A. 1952a. O polimorfismo foliar e os estudos de filometria. Aplicaçao prática de um método ampelometrico. Agron. Lusitana 4: 339–359.

Rodrigues, A. 1952b. Um metodo filométrico de caracterizaçao. Lisboa, Portugal: Fundamentos. Descripçao-Técnica Operatôria, Serviço editorial da repartiçao de estudos, informaçao e propaganda. Lisboa.

Santiago, J. L., S. Boso, J. P. Martín, J. M. Ortiz, M. C. Martínez. 2005. Characterization and identification of grapevine cultivars (Vitis vinifera L.) from northwestern Spain using microsatellite markers and ampelometric methods. Vitis 44: 67–72.

Santiago, J. L., S. Boso, P. Gago, V. Alonso-Villaverde, M. C. Martíne. 2007. Molecular and ampelographic characterisation of Vitis vinifera L.“ Albariño”,“ Savagnin Blanc” and “ Caíño Blanco” shows that they are different cultivars. Spanish Journal of Agricultural Research 5: 333–340.

Santiago, J. L., I. González, P. Gago, V. Alonso-Villaverde, S. Boso, M. C. Martínez. 2008. Identification of and relationships among a number of teinturier grapevines that expanded across Europe in the early 20th century. Australian journal of grape and wine research 14: 223–229.

Silvia, D., B. O’Shea, B. Danielak. 2019. A Learner-Centered Approach to Teaching Computational Modeling, Data Analysis, and Programming. In International Conference on Computational Science (pp. 374–388). Berlin, Germany: Springer.

Silvia, D., N. Hawkins, B. O’Shea, M. Caballero. 2020. Designing curricula for data science based on fundamental skills and competencies informed by expert interviews. Bulletin of the American Physical Society. March 4, 2020.

Wickham, H. 2016. ggplot2: elegant graphics for data analysis. Berlin, Germany: Springer.

Wilf, P., S. Zhang, S. Chikkerur, S. A. Little, S. L. Wing, T. Serre. 2016. Computer vision cracks the leaf code. Proceedings of the National Academy of Sciences 113: 3305–3310.

