## Supplementary figures and images for "Composite modeling of leaf shape across shoots discriminates *Vitis* species better than individual leaves"

### Appendix S1

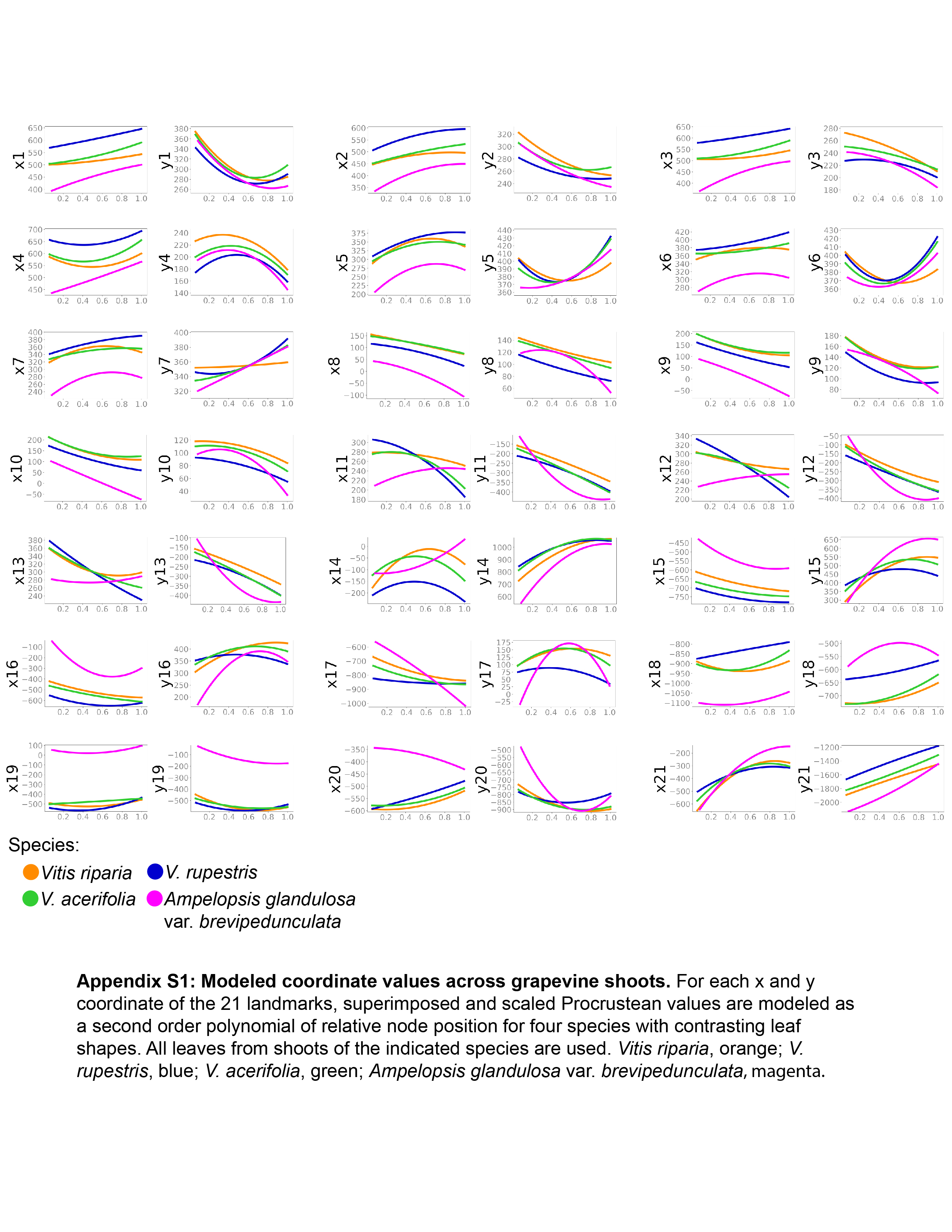
